# Hippocampal theta distinguishes between memory-guided and exploratory saccades in humans

**DOI:** 10.64898/2026.07.20.739696

**Authors:** Camilo A. Castelblanco Riveros, Peter A. Angeli, Matthijs A. A. van der Meer, Krzysztof A. Bujarski, Caroline E. Robertson

## Abstract

Memory shapes how we explore the visual world, but the neural mechanisms linking mnemonic processes to eye movements during naturalistic viewing are not well understood. Theta-band oscillatory activity in the hippocampus is time-locked with eye movements in primates, suggesting a putative mechanism for coordinating mnemonic processing and oculomotor behaviour. Yet, it remains unknown whether this coupling generalises to episodic memory-guided viewing in humans, whether slow (3–6 Hz) and fast (6–10 Hz) theta bands play dissociable functions, and whether this coupling is sensitive to the direction of upcoming eye movements. Here, we used intracranial EEG and eye-tracking data from 11 neurosurgical patients of either sex (Keles et al., 2024) to address these questions. Using fixation-locked analyses, we found that hippocampal theta dynamics differentiate memory-guided from memory-independent fixations of episodically encoded naturalistic scenes (i.e., movies) through power and coherence mechanisms that are both temporally and spectrally dissociable. First, immediately after fixations (∼0 to 250 ms), slow-theta power was more strongly suppressed during memory-guided trials (i.e., true positive, TP) than correct memory-independent trials (i.e., true negative, TN). Second, in the period around fixations (-80 to 60 ms), theta phase coherence increased independent of power during TP vs TN trials. This increase in coherence was most pronounced for contraversive relative to ipsiversive fixations, consistent with direction-sensitive hippocampal–oculomotor coordination during memory-guided viewing. Together, these findings suggest that hippocampal theta plays dissociable, time-locked roles in memory-guided fixations during naturalistic visual retrieval, supporting an ecologically relevant role for the hippocampus in coordinating memory and active visual behaviour.

**Significance statement:** Memory influences how we sample the visual environment, but the brain signals that link memory systems to eye movements are poorly understood. Using intracranial recordings from the human hippocampus during visual exploration of naturalistically encoded scenes, we demonstrate that theta-band activity differentiates memory-guided from memory-independent fixations in two ways: increased theta synchrony around fixation onset and reduced theta power modulations after fixation onset. These results reveal novel insights into how hippocampal theta helps coordinate memory-guided visual exploration.

## Introduction

Memory shapes how we visually explore the world, influencing when and where we move our eyes as we look around the world (Ryan et al., 2020). Prior work suggests that this interaction depends on coordinated hippocampal dynamics, particularly in the theta range, which are often time-locked to eye movements (Meister and Buffalo, 2016). Yet, the mechanisms by which these neural dynamics coordinate mnemonic and oculomotor processes remain poorly understood in humans, especially under naturalistic viewing conditions.

Hippocampal theta oscillations are well positioned to coordinate memory-guided visual exploration because they unfold on the timescale of saccadic eye movements. In humans, theta bouts (lasting ∼350 – 800 ms; Watrous et al., 2013) occur at a frequency of approximately 3 – 10 Hz, and are often subdivided into “slow” (3 – 6 Hz) and “fast” (6 – 10 Hz) theta bands (Lega et al., 2012; Goyal et al., 2020). Saccades occur on a similar timescale during natural visual exploration, at approximately 2.5 – 5 Hz (Wilming et al., 2017; Haskins et al., 2020). Consistent with this temporal overlap, theta bouts in primates coincide with eye movements, suggesting that theta may coordinate visual sampling with memory processing (Hoffman et al., 2013; Jutras et al., 2013; Zubair et al., 2026).

The nature of temporal coupling between theta and eye movements appears to differ across memory states. During encoding, saccades to novel stimuli reliably induce theta phase resetting and theta power (Hoffman et al., 2013; Jutras et al., 2013; Katz et al., 2020; Kragel et al., 2020), and theta power predicts subsequent recognition (Jutras et al., 2013). By contrast, during retrieval, theta phase coherence increases immediately *before* fixations to familiar (vs. unfamiliar) stimuli (Kragel et al., 2020), and theta oscillatory prevalence decreases before fixation revisits (vs. other) stimuli (Kragel et al., 2021). Thus, theta may be a mechanism by which memory prospectively guides gaze toward familiar object locations.

Here, we sought to understand how theta might prospectively guide memory-guided eye-movements during naturalistic retrieval in humans. First, whether the coupling between hippocampal theta coherence and eye movements during recall generalises beyond object–location association paradigms is unknown (Kragel et al., 2020). Does theta prospectively guide gaze during naturalistic forms of retrieval, such as scene recognition? Second, it is unclear whether slow and fast theta sub-bands differentially contribute to this phase-locking mechanism (Jacobs, 2014). Fast theta emerges from the posterior hippocampus (Goyal et al., 2020), and is associated with movement speed (Goyal et al., 2020; Maoz et al., 2023) and spatial memory retrieval (Vivekananda et al., 2020). In contrast, slow theta typically arises in the anterior hippocampus (Goyal et al., 2020) and is related to successful encoding (Lega et al., 2012) and recall (Kragel et al., 2017; Solomon et al., 2017, 2019). Whether slow vs. fast theta differentially contributes to memory-guided eye movements has not been explored. Finally, it is unclear whether theta-eye movement coupling is shaped by oculomotor planning, given evidence in the hippocampus for visual-field biases (Knapen, 2021; Silson et al., 2021; Angeli et al., 2024) and corollary discharge-like signals (Katz et al., 2022).

To address these questions, we analysed human hippocampal iEEG recordings collected during a visual recognition task using scenes that had been encoded during prior movie viewing (Keles et al., 2024). Movies provide context-rich sensory input, making them an excellent paradigm for studying more naturalistic vision and memory (Zhang et al., 2021). During recognition, participants viewed familiar scenes from the movie as well as novel scenes from the same film. We compared fixations during “memory-guided” viewing, when recently formed scene memories could guide visual sampling, with those during “memory-independent” viewing of novel scenes. By time-locking analyses to each fixation, we compared hippocampal theta power and inter-trial coherence across these two forms of visual exploration. In brief, we found that hippocampal theta dynamics differentiate memory-guided from memory-independent fixations during visual exploration of naturalistic scenes via temporally dissociable power and coherence mechanisms aligned with distinct phases of visual sampling.

## Materials and Methods

### Dataset and Paradigm

To investigate hippocampal dynamics time-locked to eye movements during a naturalistic memory recognition task, we utilised an open-access dataset consisting of intracranial EEG, eye-tracking, and behavioural data from 20 neurosurgical patients with refractory epilepsy (11 females; mean age: 37.8±13.9; all with normal or corrected-to-normal vision) who watched a short movie and subsequently participated in a visual recognition task (Keles et al., 2024; **Fig. 1A-B**). Participants’ gaze position was continuously recorded throughout the testing session using an EyeLink 1000 (SR Research Inc., Ottawa, Canada) eye-tracker (sampling rate, 500 Hz).

**Figure 1.**
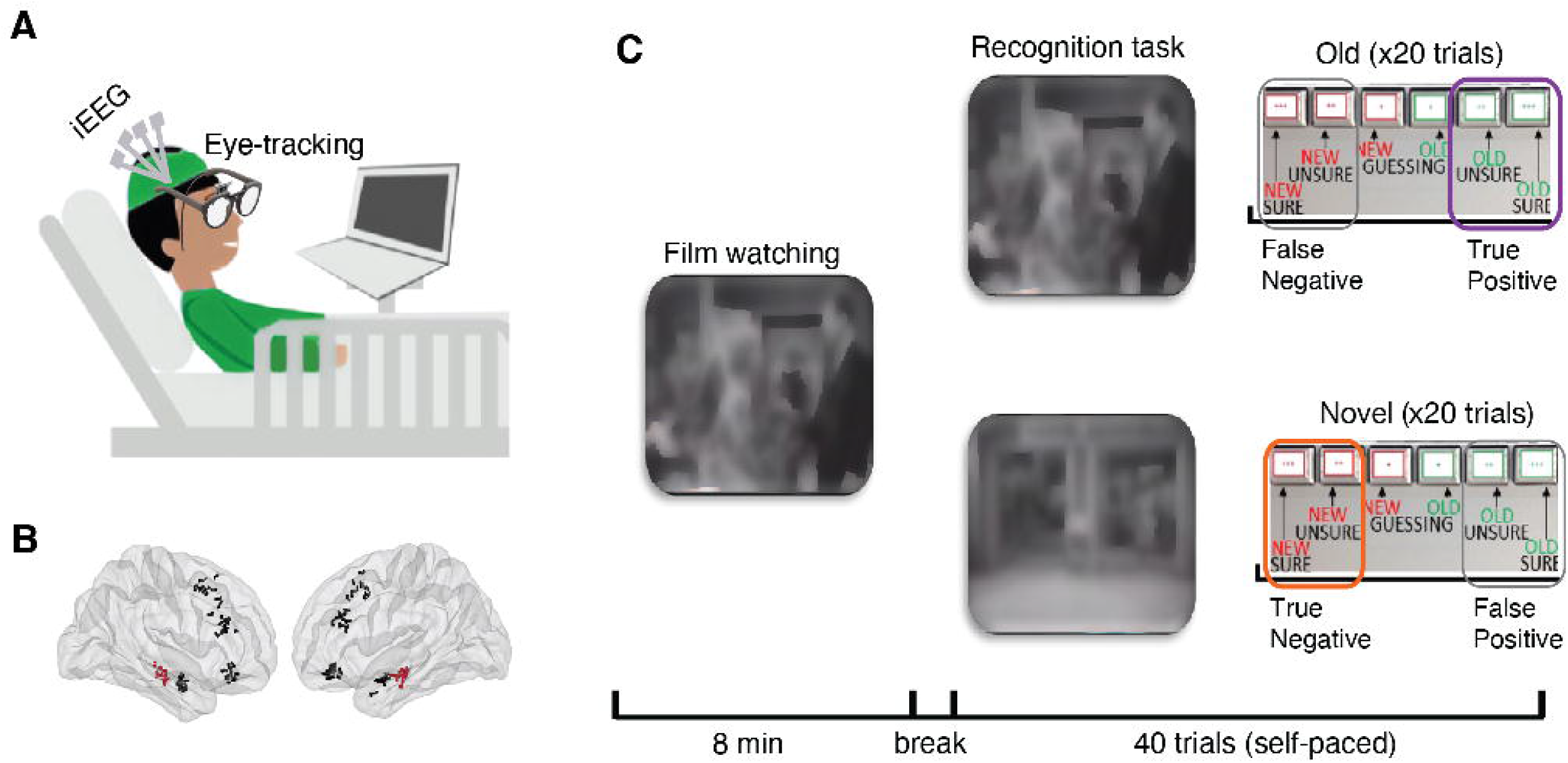
Overview of experimental paradigm and recording modalities. (A) Simultaneous intracranial EEG (iEEG) and eye-tracking recordings were obtained in neurosurgical patients. (B) Locations of electrode contacts across the 11 analysed participants, projected into MNI space. Electrodes in the shafts targeted to the hippocampus are coloured in red. (C) Experimental paradigm. Participants viewed an 8-minute film (naturalistic encoding) followed by a self-paced recognition task on 40 frames (20 old, 20 novel), yielding true-positive, true-negative, false-positive, and false-negative trials.

The testing protocol consisted of two experimental tasks: 1) a movie-watching task; 2) a scene recognition task, which assessed memory for scenes from the movie (**Fig. 1C**). These two tasks were repeated in two runs (movie watching; scene recognition; movie watching; scene recognition) to assess test-retest reliability. Since we were not interested in investigating test-retest effects, we focused solely on the recognition data from run one.

During the movie-watching task, participants watched a short (8-minute) clip from the audiovisual movie “Bang! You’re Dead” (1961). On each trial of the scene recognition task (N=40; **Fig. 1C**), participants were shown a scene (i.e., a movie frame) from the film which was either familiar (i.e., it was selected from the 8-minute segment that participants had watched during the movie viewing task; N=20) or novel (i.e., it was chosen from another point in the movie that participants had not seen before; N=20). Critically, all scenes came from the same film and thus featured similar visual content; however, only half of the scenes would have been familiar to participants based on their previous experience (i.e., the movie-watching task). On each trial, participants were asked to report whether the image was familiar or unfamiliar using a confidence rating scale that ranged from 1 (novel – sure) to 6 (familiar – sure).

Of the 16 participants for whom iEEG data were collected, we ultimately analysed data from 11 individuals (8 female, 32.73±13.96 years old) after excluding individuals with epileptic activity during the experiment (see Intracranial Recordings below) or significant missing gaze data (see Eye-tracking Data Analysis below) (Keles et al., 2024).

### Eye-tracking data analysis

#### Preprocessing

Eye-tracking data from the scene recognition task was analysed as follows. Fixation timestamps, durations, and spatial positions (x, y coordinates) were provided in the dataset (Keles et al., 2024). The first fixation in each trial was excluded to eliminate centre bias and strong image-evoked activity. Further, any fixations that lasted less than 250ms or occurred within 500ms of another fixation onset were excluded to prevent contamination by additional eye movements within the analysis window. One subject was excluded from the analysis because they had missing data for more than half of the movie-watching session.

#### Gaze patterns analysis

For each trial, we calculated mean estimates of fixation duration, count, entropy, and saccade length. Fixation duration represents the mean duration of fixations within a trial. Similarly, fixation counts represent the number of fixations made in each trial. Fixation entropy refers to the spatial entropy of fixations on each trial. To calculate fixation entropy, we first generated density maps based on duration. Then we transformed these maps into probability distribution matrices by dividing each pixel’s fixation duration by the total fixation duration across all fixations in a trial. Then, we computed Shannon’s entropy, defined as the degree of homogeneity in the probability distribution of fixations:

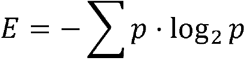

Where *p* denotes the vectorised fixation probability distribution (Açık et al., 2010; Haskins et al., 2020). Lastly, saccade length was defined as the mean Euclidean distance between all pairs of sequential fixations on the trial.

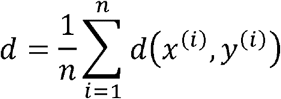

### Intracranial Recordings

#### Preprocessing

Intracranial recording procedures are described in detail in Keles et al. (Keles et al., 2024) and are reproduced briefly here. Electrophysiological data were recorded at Cedars-Sinai Medical Center using Benkhe-Fried electrodes (AdTech Inc.). Local field potential (LFP) activity was sampled at 2000 Hz (1018 electrode contacts in total; 93±20 electrodes per subject). Participants were excluded from the analysis if they had significant epileptic activity throughout the task, as determined by a board-certified neurologist. Data from hippocampal electrode contacts (hereafter referred to as ‘electrodes’) were analysed in this study.

Electrophysiological data were preprocessed using MATLAB R2024a (MathWorks Inc., Natick, MA) and FieldTrip (Oostenveld et al., 2011; Stolk et al., 2018). Continuous time-series data were first demeaned (i.e., detrended using a 0th-order polynomial) and filtered for power-line noise at 60 Hz and its harmonics (120, 180, and 240 Hz) using a zero-phase, 3rd-order Butterworth bandstop filter with a 2 Hz bandwidth (Hamming window). Data were low-pass filtered using a zero-phase, 6th-order Butterworth filter with a cutoff frequency of 475 Hz, and high-pass filtered using a zero-phase, linear-phase FIR filter of 998th order with a cutoff frequency of 0.6 Hz. All filtering was applied in a forward and reverse direction to eliminate phase distortion.

#### Analysis

Channels whose mean voltage amplitude, voltage derivative, or RMS value exceeded five standard deviations above the mean of all electrodes were flagged and subsequently excluded after visual inspection in the time-frequency domain. Channels with epileptiform activity, as determined by a certified neurologist, were also excluded. Subsequently, artefact-free data were re-referenced to a global average reference, and monopolar hippocampal derivations were selected (**Fig. 3A**). Finally, the data were epoched around fixations (±3 seconds). For epoched data, we excluded epochs with sporadic epileptic spikes by computing the signal derivative and identifying time points with absolute values exceeding 80 μV.

**Figure 2.**
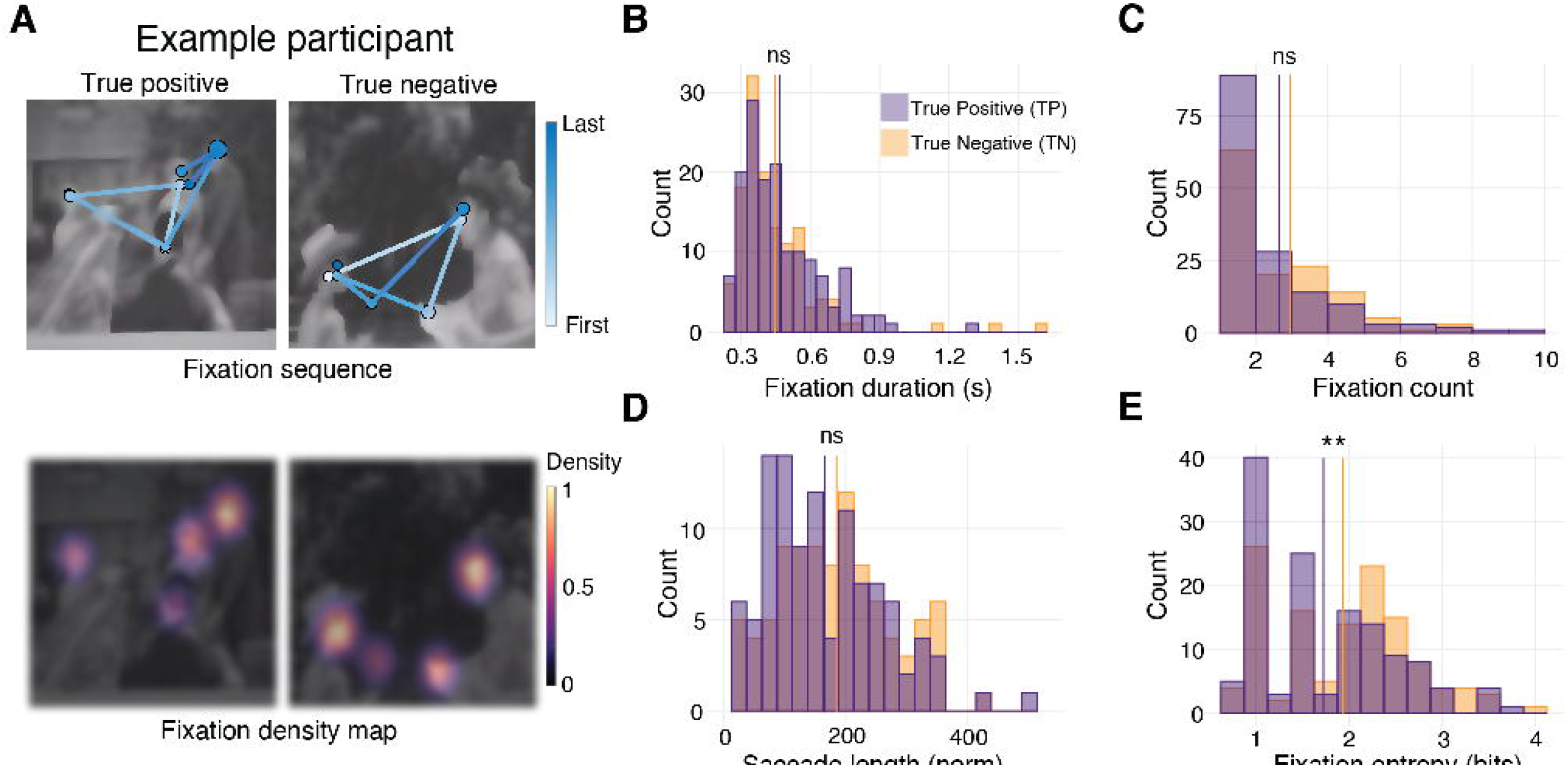
Eye movements during true-positive and true-negative trials are matched in duration and count but differ in fixation entropy. (A) Example gaze behaviour from one participant, showing fixation sequences and saccades (top) and corresponding fixation density maps (bottom) for a true positive (TP; left) and a true negative (TN; right) trial. Group-level comparisons show no significant differences between TP and TN trials in (B) mean fixation duration (*p* = 0.31), (C) fixation count (*p* = 0.10), or (D) mean saccade length (*p* = 0.13). (E) Fixation entropy was significantly larger for true negatives compared to true positives (*t*(253) = −2.51, *p* = 0.01**; β = −0.30), indicating greater spatial dispersion of gaze during memory-independent viewing. All histogram data are represented as mean values per trial across participants. \**p* < 0.05. \*\**p* < 0.01.

**Figure 3.**
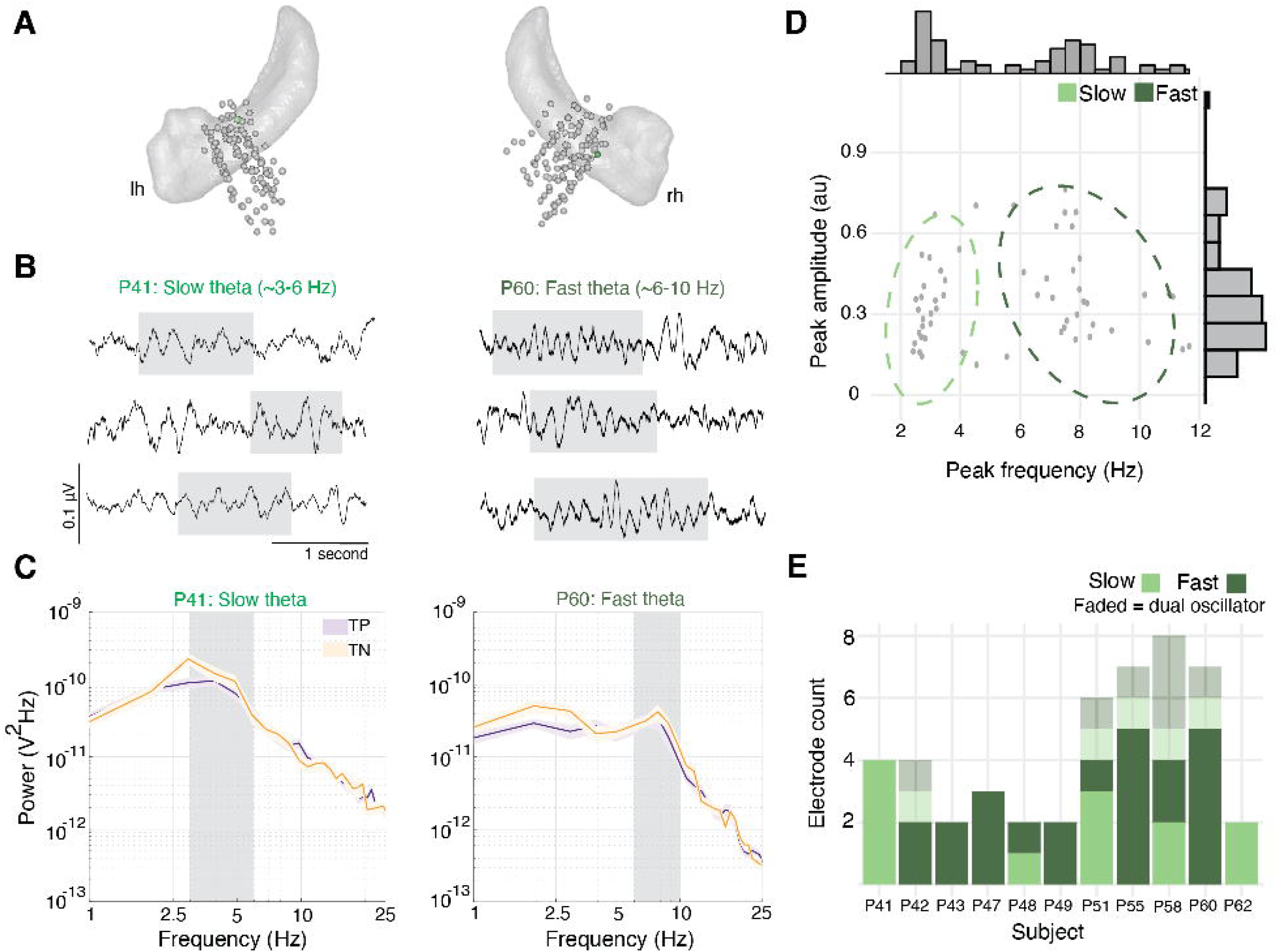
Hippocampal theta activity was robustly present during the recognition task. (A) MNI-based hippocampal electrode locations for all participants; only contacts within the hippocampus were included in analyses. (B) Example theta band traces from two representative contacts in different participants, showing slow theta (3–6 Hz; left) and fast theta (6–10 Hz; right). (C) Power spectral densities (PSDs) for the two example contacts shown in (B), plotted separately for true-positive (TP) and true-negative (TN) trials. Shaded regions indicate the theta sub-band ranges. (D) Theta peak frequency and amplitude for hippocampal electrodes with oscillatory peaks in either the slow or fast theta range. Marginal histograms show the distribution of peak frequencies (top) and amplitudes (right). (E) Number of electrodes classified as slow or fast theta per participant. Note that 8/11 subjects had both slow and fast theta electrodes. Solid bars indicate single-oscillator electrodes. Faded bars indicate dual-oscillator electrodes (N = 6), which were excluded from subsequent band-specific analyses.

### Neural power spectra

To identify hippocampal electrodes exhibiting theta activity during the memory task, we computed the power spectral density (PSD) from continuous (non-epoched) recognition task data. PSDs were estimated using multitaper spectral analysis employing Discrete Prolate Spheroidal Sequences (DPSS) tapers, yielding power estimates in 66 log-spaced frequencies ranging from 1 to 250 Hz.

To identify oscillatory activity, we utilised the FOOOF algorithm (*Fitting Oscillations and One-Over-F*, v. 1.1.0, Donoghue et al., 2020, https://fooof-tools.github.io/fooof/), which distinguishes peaks in the PSD relative to the 1/f drop-off. Electrodes with oscillatory peaks (minimum log peak power > 0.1 over 1/f) between 3 and 10 Hz were defined as theta-exhibiting electrodes and included in subsequent analyses (3.73± 1.74 electrodes per participant). As in previous reports, we identified two distinct hippocampal activity patterns at approximately 3 Hz (slow theta) and 8 Hz (fast theta).

For condition-specific analyses (TP and TN), peri-fixation data epochs (described above) were zero-padded with 1250 ms to avoid edge artefacts prior to computing PSDs or performing time-frequency analyses. Time-frequency analyses were calculated using DPSS tapers in 30 log-spaced frequencies (1 to 10 Hz). Data were baselined to the pre-fixation interval of −500 to −400 ms relative to fixation onset.

### Inter-trial phase coherence

To investigate the coupling between hippocampal oscillatory dynamics and eye movements, we computed the inter-trial phase coherence (ITC), also known as Inter-Trial Phase-Locking, around fixations (Tallon-Baudry et al., 1996; Van Diepen and Mazaheri, 2018). ITC quantifies the phase-locking consistency of oscillatory activity across repeated epochs (peri-fixations) within each electrode as:

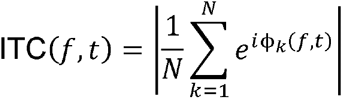

Where N is the number of observations and φ*_k_ is* the local phase angle in radians.

To compute ITC, we decomposed our 3-second iEEG epochs using a wavelet approach with 30 log-spaced frequencies between 1 and 10 Hz, a window length of 4 cycles, and computed phase estimates every 2 ms. We computed ITC for each experimental condition separately. Because the number of observations can bias ITC (Katz et al., 2020), we transformed ITC into ITC_z_ values, also known as Rayleigh’s Z, via:

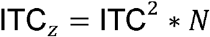

Where N is the number of observations used to compute ITC. This transformation was applied separately for each subject, electrode, and condition. Resulting ITC_z_ values were concatenated across electrodes, grouped by condition, and baseline-corrected to the pre-fixation interval (−500 to −400 ms) prior to statistical analyses.

### Data Whitening

To explicitly control for the 1/f trend in iEEG data and distinguish between oscillatory changes and broadband activity, we repeated key power and coherence analyses after performing temporal whitening (**Fig. S3, S4, and S7**) by taking the difference between consecutive time points prior to computing power and ITC estimates (Groppe et al., 2013; Kleinfeld and Mitra, 2014).

**Figure 4.**
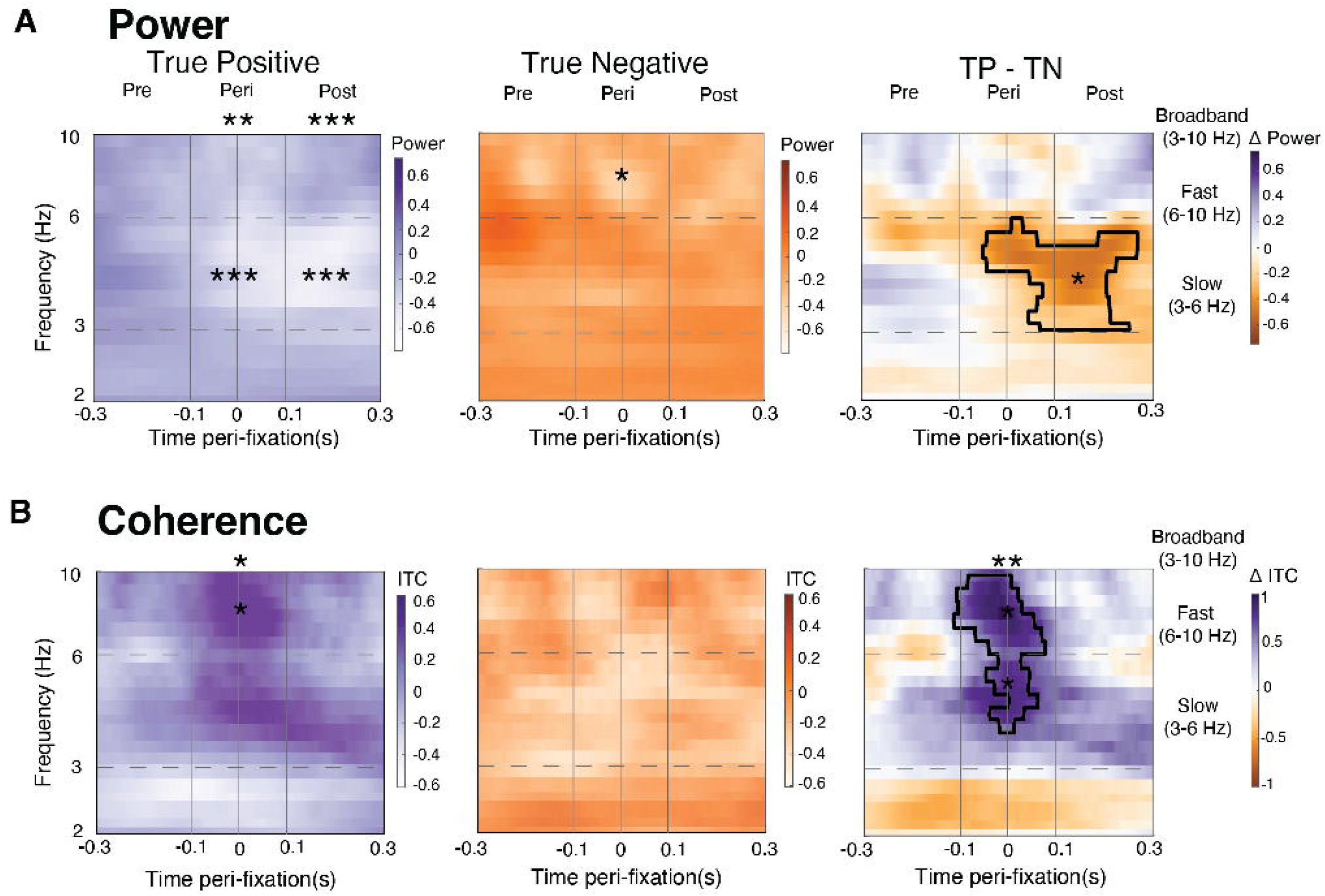
True positive and true negative trials show dissociable peri-fixation signatures in both power and coherence. (A) Time-frequency power spectrograms peri-fixation for TP (Left) and TN (Middle) trials, averaged across all theta electrodes. Direct contrasts between conditions (TP − TN; right) revealed lower power for TP than TN trials, beginning around fixation onset and extending into the post-fixation period, with the strongest effect in the slow-theta range. (B) Inter-trial coherence (ITC**_z_**) aligned to fixation onset for TP (Left) and TN (Middle) trials. Direct contrasts between conditions (TP − TN; right) revealed greater peri-fixation phase coherence for TP vs. TN trials, spanning approximately −80 to 60 ms relative to fixation onset. Dashed horizontal grey lines denote 3 and 6 Hz. Vertical lines (-100, 0, and 100ms) denote pre-, peri-, post-fixation time periods. Asterisks indicate significant differences from baseline within the pre-specified slow-, fast-, or broadband-theta windows (*p<0.05, **p<0.01). Black outlines indicate periods of significant differences based on non-parametric permutation testing.

### Event-Related Potentials

To assess whether ITC estimates were driven by event-related potential (ERP) artefacts rather than true phase-clustering (**Fig. S8**), we computed fixation-onset ERPs for each theta electrode by averaging iEEG data within a broad window (±1.5 s around fixation onset) across all fixation epochs, then repeated our ITC analysis after subtracting this average ERP (without baseline correction).

### Statistical analyses

Participants rated images as either novel or old using a 6-point confidence scale, from 1 (novel, sure), 3 (novel, guessing), 4 (old, guessing) to 6 (old, sure). Based on these ratings and trial status, we classified trials into: Correctly remembered (true positive, TP), correctly identified as novel (true negative, TN), incorrectly remembered (false positive, FP), and incorrectly identified as novel (false negative, FN). To reduce behavioural noise, we included only trials with high-confidence responses, specifically ratings 1–2 (high-confidence “novel”) and 5–6 (high-confidence “old”).

To compare eye-tracking metrics between conditions (e.g., fixation count, saccade length, etc.), we used linear mixed-effects models to account for non-independence of observations within subjects and for unequal numbers of fixations across subjects and conditions. This approach allowed us to estimate condition effects while controlling for inter-individual variability. Four metrics (mean fixation, count, entropy, and saccade length) were modelled independently as:

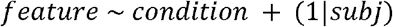

Time-frequency power and coherence metrics (ITC_z_) were compared across conditions using two methods: non-parametric permutation testing and linear models. Non-parametric permutation testing was used to control for multiple comparisons across both time and frequency dimensions (Maris and Oostenveld, 2007). Condition labels across electrodes were randomly permuted 5,000 times to generate null distributions and establish cluster-forming thresholds. Clusters were formed by grouping adjacent time-frequency points exceeding a cluster-forming threshold (alpha = 0.05). The statistical significance of each cluster was assessed by comparing the observed cluster-level t-values to the null distribution derived from permutations, using a two-tailed threshold of p < 0.025 to control for the family-wise error rate. The analysis focused on the interval from −300 ms to 300 ms and frequencies between 3 and 10 Hz. Effect sizes were computed by averaging the metric of interest (i.e., power or coherence) within significant clusters for each unit of observation, collapsing across time-frequency dimensions. These per-unit cluster-averaged values were subsequently used to compute group-level effect sizes, expressed as Cohen’s d.

Linear models were used as a complementary confirmatory analysis to quantify condition effects within predefined theta bands and peri-fixation time windows. Specifically, we compared mean power and coherence estimates across conditions at −100 to 100 ms (peri), −250 to −50 ms (pre), and 50 to 250 ms (post) for slow, fast, and broadband theta frequencies, using the following model: *feature ∼ condition.* We also examined whether electrodes exhibited differential responses across conditions and theta bands by fitting the following interaction model to each time window: *feature ∼ condition * theta band*. Additionally, for each condition independently, we assessed whether mean coherence and power were significantly different from zero using one-tailed t-tests. Finally, we compared coherence between contraversive and ipsiversive fixations using the following model: *feature ∼ condition*direction*.

## Results

### Gaze behaviour reflects memory-guided viewing

On each trial of a visual recognition task (N=40), participants viewed a greyscale scene depicting either a familiar frame from a movie they had watched (“old”; n=20) or a novel frame from the same film (“novel”; n=20) (**Fig. 1**). Overall memory performance was high, with an 85.00, ± 11.62% hit rate (mean ± sd), 70.91, ± 12.41% correct rejection rate, and 77.95 ± 8.20% accuracy. On average, individuals had 17.0 ± 2.3 trials in which scenes were correctly recognised as old (true positives, TP; 42.5% ± 5.8% of total trials) and 14.2 ± 2.5 trials in which scenes were correctly recognised as old (true negatives, TN; 35.5% ± 6.2% of total trials) (**Fig. S1A–B**). This high task performance motivated us to focus our primary analyses on TP and TN conditions, which differed only in memory availability, while being matched for response accuracy and trial numbers. Specifically, TP trials involved correctly recognised familiar scenes, for which participants could use recently formed scene memories to guide familiarity judgments and visual sampling. In contrast, TN trials involved correctly rejected novel scenes, for which no scene-specific memory was available.

Across subjects and conditions, a total of 2402 of fixations occurred; the number of fixations was comparable for TP and TN trials (t = 0.482, p = 0.63; β = 0.04). Of all fixations, 42.58% were removed because they lasted less than 250ms. Previous work has revealed that during active viewing behaviours, fixations tend to last around 250ms (Haskins et al., 2020). To prevent overlap between two subsequent fixations during the time window of interest, we also excluded any fixation that occurred within 500ms of a previous fixation (14.40% of total fixations). Thus, a total of 1036 fixations were included in the main analyses (subject mean TP: 36.18 ± 12.84 STD; TN: 34.36 ± 12.52 STD).

Next, we quantified fixation duration, fixation counts, fixation entropy, and saccade length for TP and TN trials (**Table 1**, **Fig. S1C**). Linear mixed-effect modelling showed that TP and TN trials were matched for fixation duration (*t*(276) = 1.01, *p* = 0.31; β = 0.12; **Fig. 2B**), fixation count (*t*(276) = −1.66, *p* = 0.10; β = −0.19; **Fig. 2C**), and saccade length (*t*(201) = −1.51, *p* = 0.13, β = −0.21; **Fig. 2D**). In contrast, fixation entropy was smaller in TP compared to TN trials, indicating less spatially distributed and less exploratory gaze patterns when observing familiar stimuli (*t*(253) = −2.51, *p* = 0.01; β = −0.30; **Fig. 2E**), consistent with prior findings (Smith and Squire, 2008; Althoff and Cohen, 1999). Thus, TP and TN trials isolate memory-related differences in neural activity during fixation, independent of differences in fixation duration, fixation count, or saccade length.

#### Hippocampal theta activity is robustly present during the memory recognition task

Having established that TP and TN trials were closely matched in oculomotor behaviour and trial counts, we next examined whether hippocampal theta activity differed during the period surrounding fixations in memory-guided (TP) vs. memory-independent (TN) trials. We first identified hippocampal electrodes that exhibited theta-band activity (3 −10 Hz) during recognition task trials (**Fig. 3A**). For each electrode, we computed power spectral densities (PSDs) across all task trials. We then used the FOOOF algorithm to parameterise each power spectrum, separating oscillatory peaks from the aperiodic 1/f background (Donoghue et al., 2020). Electrodes with significant theta peaks between 3 and 10 Hz were designated as “theta electrodes” and used for all subsequent analyses (41/63 electrodes; 65.10%; see **Fig. 3A-D**).

Across electrodes, theta peaks tended to cluster around ∼3 and ∼8 Hz, consistent with previous descriptions of slow/low and fast/high theta, respectively (Goyal et al., 2020) (**Fig. 3C**). Most participants had electrodes exhibiting both slow- and fast-theta activity (8/11 participants; **Fig. 3E**). Because electrode coverage across participants was primarily confined to the mid-portion of the hippocampus (**Fig. 3A**), analyses comparing anterior versus posterior hippocampus were not feasible.

Overall, hippocampal theta-band power was comparable between TP and TN trials in both the slow and fast theta bands across all electrodes (both *p* > 0.05) (**Fig. 3C**). Visual inspection of hippocampal traces revealed that both fast and slow theta-band oscillatory activity tended to occur in intermittent bouts (<u>Jutras et al., 2013</u>), unlike the ongoing, continuous theta-band oscillations seen in rodents (<u>Buzsaki, 2002</u>) **(Fig. 3B)**. Theta-band oscillatory activity (i.e., theta bouts) was identified during task trials using the BOSC toolbox (<u>Kosciessa et al., 2020</u>; <u>Whitten et al., 2011</u>) to detect oscillatory events lasting at least three cycles (**Fig. S2A**). The mean theta *P_episode_*(i.e., proportion of time with BOSC-detected oscillations over total trial time) was 0.166 ± 0.082, comparable with previous studies in primates (<u>Jutras et al., 2013</u>) and humans (<u>Ekstrom et al.,</u> <u>2005</u>). No differences in *P_episode_* were observed between TP and TN conditions (*p* > 0.05). Thus, theta activity was robustly detected during both TP and TN trials.

#### Theta power is reduced after fixation onset

Given the robust presence of hippocampal theta activity during the recognition task, we next asked whether theta power was modulated in the time window around fixation onset (±300 ms) and whether this modulation differed between TP and TN trials (**Fig. 4A**). Spectral power was baseline-normalised to a pre-fixation window (-500 to -400ms), which did not differ between conditions (cluster P_FWE_ > 0.05). Non-parametric permutation testing across the peri-fixation interval (−300 to 300 ms; 3–10 Hz) revealed a significant TP–TN difference, with lower power for TP than TN trials from approximately fixation onset into the post-fixation window, primarily within the slow-theta range (∼3-6 Hz; cluster P_FWE_ = 0.017; d = −0.580; **Fig. 4A**). This TP–TN difference was also evident without baseline normalisation, with subtraction maps showing greater post-fixation theta power for TN than TP trials (p<0.05) (**Supp 3A**).

To identify which theta sub-bands (slow, fast, broad) and fixation-centred temporal epochs (pre, peri, post) drove the condition effect (TP vs. TN), we used linear models applied to activity summarised within predefined pre (−250 to −50 ms), peri (−100 to 100 ms), and post fixation (50 to 250 ms) windows (**Table 2**; **Fig. 4A**). First, examining each condition relative to baseline, TP trials showed significant power reductions within the slow theta range (∼3-6 Hz) during both peri- (β = -0.336 ± 0.086, p < 0.001) and post-fixation windows (β = -0.433 ± 0.091, p < 0.001), whereas TN trials did not (all p>0.05). Second, direct TP–TN comparisons showed no condition difference before fixation onset, but revealed lower power in slow theta for TP vs. TN after fixation onset, reaching significance in the post-fixation window (β = -0.335 ± 0.144, p = 0.023 Importantly, while post-fixation TP–TN power differences were primarily observed in the slow-theta band, formal interaction testing found only a marginally significant condition × theta-band effect (p = 0.09). Altogether, we observed a post-fixation reduction in theta power in TP versus TN trials, qualitatively centred in the slow theta subband.

To clarify whether these results reflected oscillatory changes rather than broadband activity, we temporally whitened the data before computing spectral power estimates (Kleinfeld and Mitra, 2014), thereby controlling for the 1/f trend in iEEG data (Groppe et al., 2013). After whitening, TP trials continued to show reduced power during the post-fixation window, and direct TP–TN contrasts again revealed lower post-fixation power within slow theta for TP than TN trials (p=0.05) (**Fig. S4A**). No differences were observed when testing for a significant condition × theta-band effect (p = 0.14). In addition to temporally whitening our data, we analysed the ∼20% of trials where theta bouts were detected (**Fig. S2B**). In this restricted and underpowered analysis, we observed that theta power post-fixation did not differ across conditions but showed the same directionality as stated earlier: greater TN power post-fixation relative to TP trials. Because this difference was not statistically significant, we cannot rule out the possibility that aperiodic changes within the theta range contributed to these power differences.

Supplementary analyses of False Positive trials (in which participants incorrectly indicate having previously seen novel images) showed no power effects in slow theta post-fixation (**Fig. S5A**). Thus, the observed slow-theta decrease was specific to memory-guided fixations and did not simply reflect participants’ subjective feeling of familiarity.

#### Increased Theta Synchrony Around Memory-Guided Fixations

Next, we tested whether theta coherence was coupled with eye movements and whether this coupling differed between memory-dependent and memory-independent trials. To test this, we first computed inter-trial phase coherence (ITC_z_; see Methods) for each electrode peri-fixation (± 300ms) (see example traces in **Fig. S6**). As for the power analyses, ITC was baseline-normalised to a pre-fixation window (-500 to -400ms), which did not differ between TP and TN trials (cluster P_FWE_ > 0.05). Non-parametric permutation testing revealed greater peri-fixation coherence for TP relative to TN trials, from approximately –80 ms to 60 ms, spanning both slow and fast theta sub-bands (∼4 – 9 Hz; cluster P_FWE_ = 0.024; *d* = 0.715; **Fig. 4B**). These phase clustering results were spectrally confined to theta frequencies (**Fig. S7**) and remained significant when coherence estimates were not baseline normalised, arguing for their robustness (**Fig. S3B**).

As described previously for theta power, we fit a series of linear models for each sub-band and epoch to characterise the TP vs. TN effect (**Table 3**; **Fig. 4B**). First, examining each condition relative to baseline, TP trials showed significant broadband theta ITC increases during the peri-fixation window (β = 0.431 ± 0.274, *p* = 0.017), whereas TN trials remained at baseline across theta bands (*p* > 0.05). Second, direct TP–TN comparisons showed greater theta coherence for TP than TN trials during the peri-fixation window, with significant effects in slow, fast, and broadband theta (all p < 0.05). No significant TP–TN differences were observed in the pre- or post-fixation epochs (all p > 0.05). Third, formal condition × theta-band interactions were not significant, indicating no evidence that the coherence effect differed across slow and fast theta bands (p > 0.05). Overall, these results indicate that a peri-fixation increase in theta coherence is selective to memory-guided fixations.

It is unlikely that these peri-fixation differences between TP and TN trials were driven by oculomotor artefacts. Fixation duration, count, and length were matched across conditions (**Fig. 2. B−D**), and retinal repositioning artefacts should affect memory-guided and memory-independent fixations similarly. Further, electrooculogram (EOG)-related artefacts increase power in the ∼20-200 Hz range and typically last ∼20ms (Yuval-Greenberg et al., 2008; Jerbi et al., 2009; Kovach et al., 2011), whereas the effect observed here is a sustained theta-band coherence difference around fixation onset. Nevertheless, we conducted an additional analysis to account for this possibility by subtracting event-related potentials (ERPs) from each channel prior to computing ITC. The peri-fixation theta coherence effect remained significant after ERP subtraction, demonstrating our findings are not driven by EOG-related artefacts (**Fig. S8**).

These phase clustering results also remained qualitatively similar when different numbers of wavelet cycles were used to compute ITC and when TP coherence was compared against False Positive trials (**Fig. S5B**). Moreover, these results differed temporally from the power results (see *Theta power is reduced for after fixation onset* above), suggesting ITC reflects a distinct neural process (Van Diepen and Mazaheri, 2018), not confounded by amplitude changes. Coherence increased peri-fixation (−80 to 60 ms) in theta broadly, whereas power decreased post-fixation (0 to 250 ms). To examine whether these coherence results reflected oscillatory differences or aperiodic modulations, we restricted our analyses to the ∼20% of trials exhibiting significant theta-activity bouts (**Fig. S2C**). While statistically underpowered, we observed a similar qualitative directionality of the main effects: increased TP coherence peri-fixation relative to TN trials. However, because these results did not survive cluster-based permutation testing, they cannot fully exclude an aperiodic contribution.

As with spectral power, we performed additional analyses in which data were whitened prior to computing ITC estimates to remove broadband contributions from oscillatory processes. Linear modelling of ITC estimates supported the temporal specificity of our main results, revealing that TP ITC was significantly greater than TN during the peri-fixation window across broadband theta (**Fig. S4B**). Together, these findings indicate that memory-related modulation of hippocampal theta is expressed through distinct changes in power and phase coherence around fixations, consistent with a mechanism in which theta activity coordinates memory-dependent perceptual sampling.

#### Slow and fast theta electrodes show similar peri-fixation trends

Next, we explored whether the power and coherence results were differentially expressed in the electrodes with slow-versus fast-theta peaks. For this electrode-specific analysis, we classified theta electrodes according to their spectral peaks. Electrodes with a single oscillatory peak in the 3–6 Hz range were classified as slow-theta electrodes (N = 12), whereas electrodes with a single peak in the 6–10 Hz range were classified as fast-theta electrodes (N = 23). Electrodes with peaks in both the 3–6 Hz and 6–10 Hz ranges, separated by at least 0.5 Hz, were classified as dual oscillators (N = 6) and excluded from this electrode-specific analyses (Goyal et al., 2020). Most participants had both slow- and fast-theta electrodes (8/11 participants; **Fig. 3E**).

To separate oscillatory activity from aperiodic 1/f contributions, we temporally whitened the data before estimating fixation-locked power and coherence (Groppe et al., 2013; Kleinfeld and Mitra, 2014). Peri- and post-fixation power was moderately lower in TP trials relative to TN trials in both slow- and fast-theta electrodes; this effect was numerically consistent with the main analysis but did not reach statistical significance in either electrode type (**Fig. 5A**). Pre- and peri-fixation coherence was greater in TP trials than TN trials in both slow- and fast-theta electrodes; this effect reached significance in fast-theta electrodes (p<0.01) but did not reach significance in slow-theta electrodes (**Fig. 5B**). Given the reduced sample size after separating electrodes by spectral profile, these results should be interpreted cautiously. Overall, however, the direction of effects was numerically consistent across electrode types, suggesting that the main peri-fixation effects were not driven exclusively by either slow- or fast-theta electrodes.

**Figure 5.**
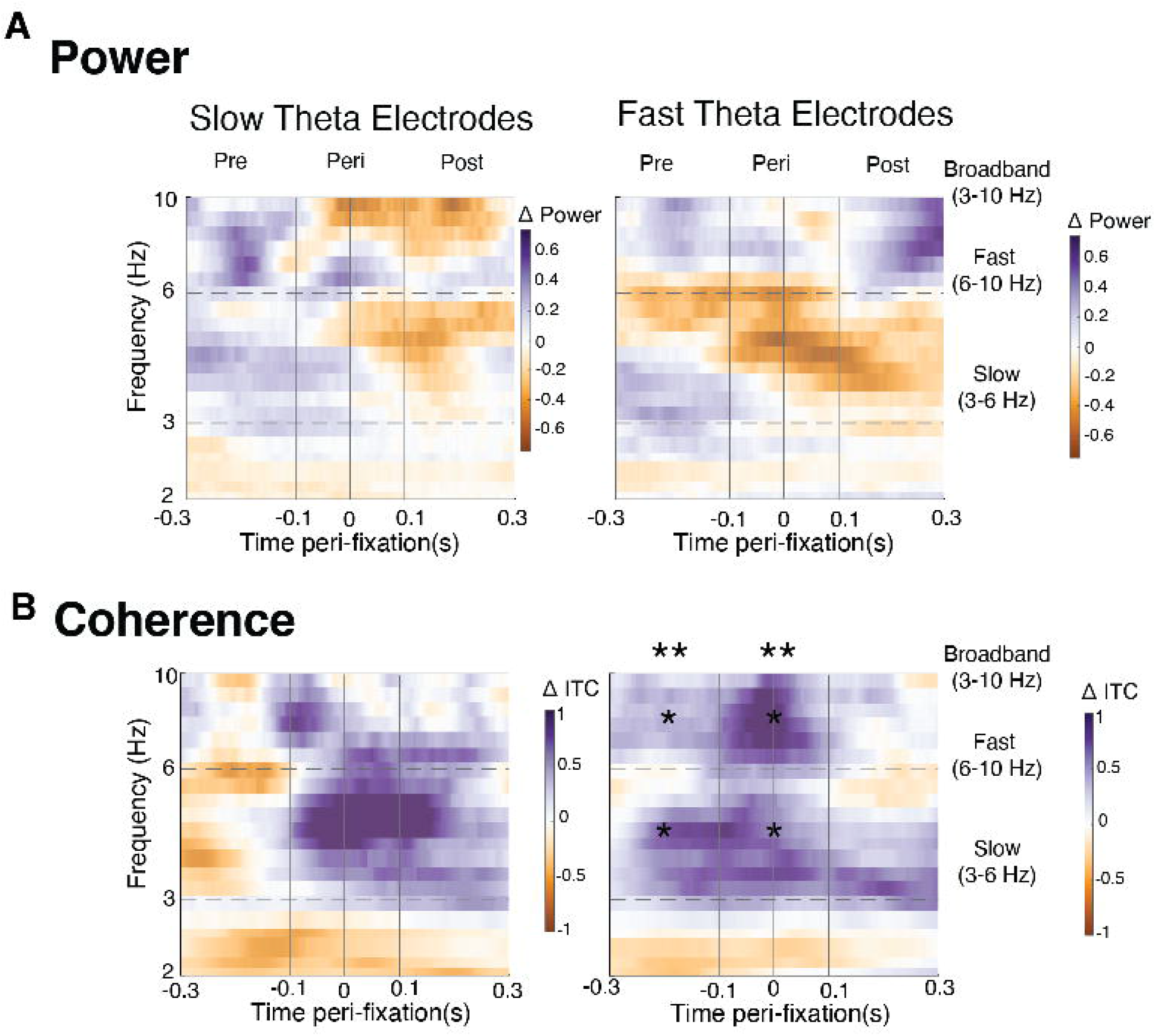
Effects across slow- and fast-theta electrodes. (A) Time-frequency power spectrograms aligned to fixation onset for slow (Left) and fast (Right) theta-electrodes, depicting direct contrasts between TP and TN trials. Across electrode types, TP trials exhibited lower spectral power peri- and post-fixation than TN trials, but the difference did not reach statistical significance. (B) Inter-trial coherence (ITC_z_) aligned to fixation onset was greater for TP trials across electrode types and, based on linear modelling, did not reach statistical significance in the pre- and peri-fixation periods for slow-electrodes (Left), but did for fast-theta electrodes (Right). Dashed horizontal grey lines denote 3 and 6 Hz. Vertical lines (-100, 0, and 100ms) denote pre-, peri-, post-fixation time periods. Asterisks indicate significant TP–TN differences from linear models within predefined theta bands and time windows (*p<0.05, **p<0.01).

#### Directional Modulation of Theta Coherence during Memory-guided Saccades

Given the observed increase in theta coherence around memory-guided fixations, we next tested whether this phase synchrony was modulated by saccade direction (i.e., left-vs. right saccades) – motivated by recent reports of directionally-tuned corollary discharge-like suppression of neural activity around saccades in mnemonic structures (Katz et al., 2022) and visual field biases in the hippocampus (Knapen, 2021; Silson et al., 2021; Angeli et al., 2024). Specifically, we tested whether ITC_z_ differed based on the direction of an upcoming saccade during TP and TN trials.

To do so, we labelled fixations as ipsiversive (saccades toward the same side as the recording electrode; e.g., rightward saccades for a right-hemisphere electrode) or contraversive (saccades toward the opposite side; e.g., leftward saccades for a right-hemisphere electrode). Using non-parametric permutation testing, we observed that theta coherence was greater in TP compared to TN trials during contraversive, but not ipsiversive, fixations in the theta range between –80 to 300ms around fixation (P_FWE_ = 0.012; *d* = 0.842; **Fig. 6**). Directional effects were specific to coherence, with no corresponding modulation of theta power (**Fig. S9**). Furthermore, the proportion of saccades that crossed the centre was comparable across conditions, at around 11% of their corresponding total (*t* = 0.742, *p* = 0.46; β = 0.02). Therefore, these ITC results cannot be explained by differences in centre-crossing saccades.

**Figure 6.**
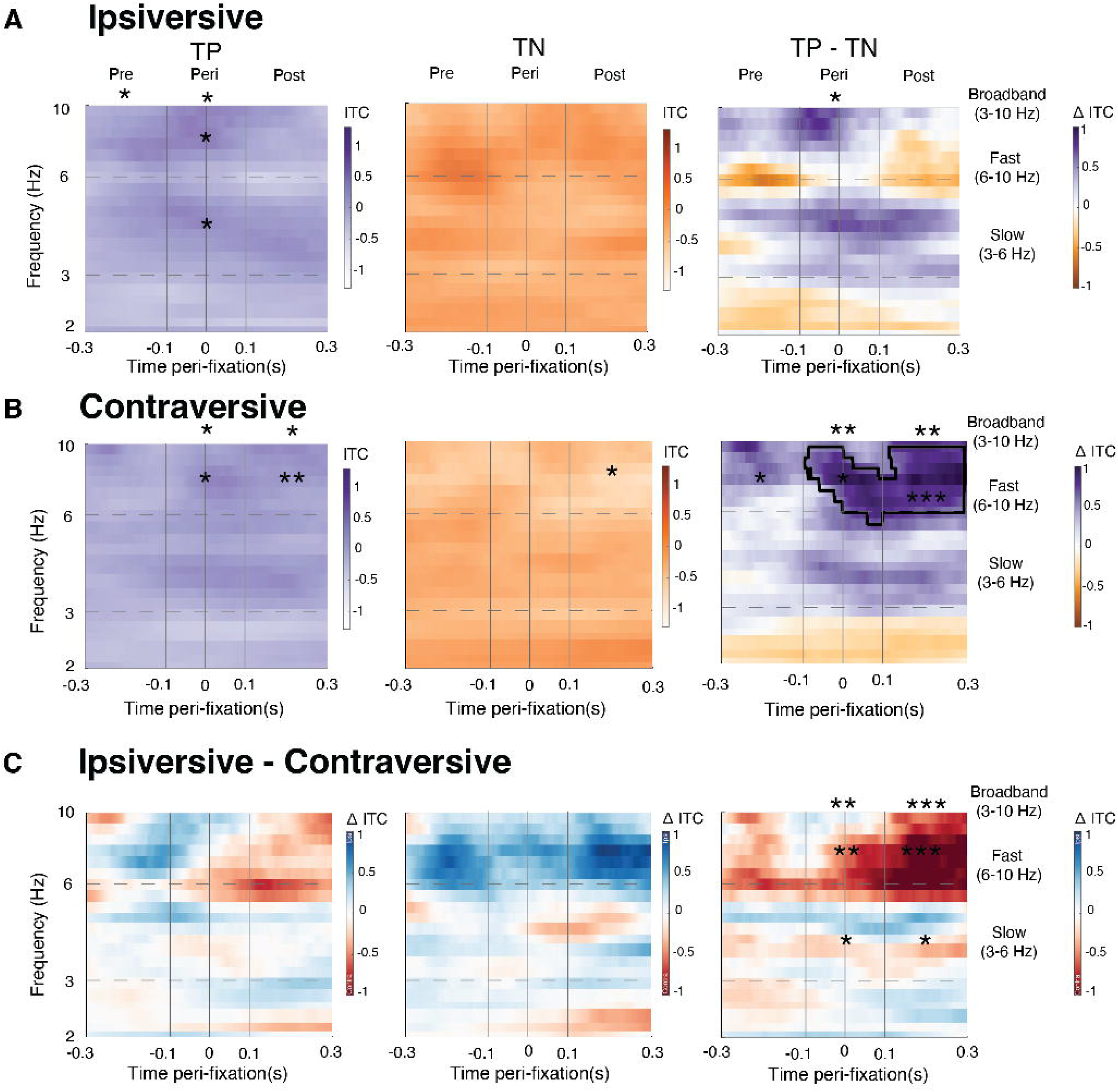
**True positive and true negative trials differ only in coherence around contraversive fixations**. (A-B) Peri-fixation inter-trial phase coherence (ITC_z_) maps showing mean phase-locking consistency for true positive (TP; Left), true negative (TN; Middle) trials, and subtraction maps across conditions (TP – TN; Right), separated by ipsilateral (A) and contralateral (B) fixations. Greater phase coherence was observed for contraversive TP trials from approximately -80 to 300 ms relative to fixation onset (P_FWE_ = 0.012, *d* = 0.842, nonparametric cluster corrected). (C) Within-condition subtraction maps between ipsiversive and contraversive ITC_z_ maps for TP (Left), TN (Middle), and TP – TN (Right) trials. Blue colours indicate more ipsiversive ITC_z_. Red colours indicate more contraversive ITC_z_. Dashed horizontal grey lines denote 3 and 6 Hz. Vertical lines (-100, 0, and 100ms) denote pre-, peri-, post-fixation time periods. Asterisks indicate statistical significance for linear models comparing TP and TN trials (*p<0.05, **p<0.01). Black outlines indicate periods of significant differences based on non-parametric permutation testing.

Finally, to characterise the dynamics of these peri and post-fixation results, we fit linear models for each sub-band and epoch as described above. In contraversive fixations, theta coherence increased for TP fixations and decreased for TN fixations during the post-fixation period (TP: β = 0.359 ± 0.108, *p* = 0.002; TN: β = −0.408 ± 0.180, *p* = 0.029; **Table 4**). During the same post-fixation period, ipsiversive fixations exhibited broadband theta coherence that remained at baseline (*p* > 0.05) (**Table 4**). In line with these results and overall directional selectivity, we observed that the TP–TN difference was greater during contraversive relative to ipsiversive fixations (β = −0.763 ± 0.294, *p* = 0.010; **Table 4**). Taken together, these results suggest that theta-band synchrony is directionally modulated during memory-guided fixations, potentially reflecting the integration of oculomotor and mnemonic signals during active viewing.

## Discussion

Memory and eye movements are time-locked during active vision, but the mechanisms through which memory systems guide oculomotor behaviour remain poorly understood. Here, we identify two dissociable hippocampal theta mechanisms that distinguish memory-guided from memory-independent visual sampling during episodic retrieval following naturalistic movie encoding (Keles et al., 2024). Memory-guided fixations were associated with a post-fixation suppression of theta power, consistent with enhanced mnemonic processing after memory-guided sampling. Additionally, we observed a peri-fixation increase in broadband theta phase coherence consistent with a modulatory input relayed to mnemonic centres. This coherence effect was direction-sensitive, with stronger coupling for contraversive fixations relative to the recorded hippocampus, revealing lateralised coordination between mnemonic and oculomotor systems.

We use “memory-guided” to refer to visual sampling during familiar scenes, where recently formed memory representations are retrieved to guide ongoing exploration. Consistent with prior work showing that prior experience reduces exploratory sampling of familiar scenes (Kaspar and König, 2011; Chakravarthula et al., 2025), fixation entropy was lower during memory-guided trials, indicating a more spatially structured gaze. This behavioural result suggests that recently formed scene memories reduced exploratory sampling and promoted selective, memory-constrained visual exploration, providing a behavioural complement to the neural effects.

Previous work linking hippocampal theta coherence to eye movements during memory retrieval has primarily relied on associative object–location paradigms, in which participants learn the spatial locations of target objects within scenes and subsequently search for them during retrieval. Within these tasks, theta phase coherence increases prior to fixations directed toward familiar object locations (Kragel et al., 2020), and theta prevalence decreases before fixation revisitations (Kragel et al., 2021). These observations suggest theta phase coherence facilitates memory-guided fixations. Still, these observations are embedded within discrete, goal-directed tasks with explicit search targets and constrained viewing behaviour, leaving unresolved how theta dynamics unfold during episodic and naturalistic settings. In the context of ambulatory spatial navigation, theta phase resetting accompanies saccades regardless of memory demand (Zubair et al., 2026). Building on these findings, we show that theta coherence increases selectively for memory-guided fixations during unconstrained, naturalistic episodic retrieval, peaking shortly before fixation onset (approximately –80 to 60 ms).

This increase in coherence is direction-sensitive, with stronger coupling for contraversive fixations, linking hippocampal theta to the motor context of ongoing visual sampling. However, we cannot fully exclude aperiodic contributions to these findings.

A separate body of work has characterised theta power dynamics during memory retrieval, largely independent of eye-movement measurements. Decreases in theta-band power have been associated with successful retrieval states (Burke et al., 2014; Kragel et al., 2017; Solomon et al., 2017, 2019; Fellner et al., 2019; Weidemann et al., 2019) and linked to increased information processing demands and attentional engagement (Hanslmayr et al., 2012, 2016; Herweg et al., 2020). However, because these studies were not fixation- or saccade-locked, the temporal relationship between theta power modulation and visual sampling has remained unresolved. Importantly, recent work has implicated theta power (5-8 Hz) in memory-dependent eye movements in ambulatory humans, showing that medial temporal lobe theta power selectively increases during saccades in memory-guided navigation compared with visually guided navigation (Zubair et al., 2026). Using fixation-locked analyses, our work shows that hippocampal theta-band power is suppressed following memory-guided fixations, suggesting that – in contrast to theta coherence effects which prospectively guide visual sampling – theta power effects might reflect post-saccadic mnemonic processing.

These effects are constrained to the theta range and tend to fall within the slow 3-6 Hz range, but cannot be robustly attributed to a specific oscillatory band. Further, the possibility that aperiodic changes within the theta range influenced these results cannot be excluded.

The temporal dissociation between theta power suppression and broadband theta coherence argues against a power-driven phase-reset account (Van Diepen and Mazaheri, 2018). Instead, it supports the interpretation that power and coherence index distinct underlying neural processes. Namely, post-fixation theta power suppression is associated with mnemonic processing, while peri-fixation theta synchrony is well positioned to organise hippocampal activity in anticipation of upcoming, memory-relevant perceptual sampling. Importantly, these results do not establish causal directionality between hippocampal activity and eye movements; rather, they reveal temporally precise signatures of coordination during naturalistic episodic retrieval.

The peri-fixation timing and direction-selectivity of theta coherence could be consistent with a corollary discharge-like (CD) mechanism linking oculomotor and hippocampal circuitry. In this frame, transient inhibition of the mesial temporal lobe *after* saccade onset resets the hippocampal theta phase, preparing pyramidal cells to process incoming visual information (Doucet et al., 2020; Katz et al., 2022). Notably, the direction-selective coherence effect — stronger for contraversive than ipsiversive fixations — was not accompanied by a corresponding power difference, suggesting that CD-like inhibition reorganises theta phase without producing a detectable amplitude signature. Functionally, CDs are thought to encode the directionality of ensuing saccades to maintain stable visual percepts (Wurtz, 2018). Mechanistically, saccade direction modulates low-frequency phase dynamics and the activity of individual neurons, potentially to filter competing inputs (Sobotka et al., 1997; Killian et al., 2015; Doucet et al., 2020; Katz et al., 2022). In humans, a CD signal may serve to create conceptual maps (Buzsáki and Moser, 2013). However, given the inherent variability of naturalistic exploration, more refined single-unit or causal approaches will be needed to determine how directional corollary discharge signals shape hippocampal theta during memory-guided viewing.

More broadly, these results reinforce a growing body of evidence demonstrating a close functional coupling between the hippocampus and visual systems (Nau et al., 2018; Turk-Browne, 2019). From this perspective, the hippocampus is not merely a recipient of visual information, but an active component of a perception-action loop that shapes how visual information is sampled. Work in non-human primates has shown that hippocampal and medial temporal lobe activity is tightly coupled to saccade timing and visual exploration, consistent with the idea that memory systems actively structure sensory sampling rather than passively encoding its outcomes (Jutras and Buffalo, 2010; Buffalo et al., 2011; Meister and Buffalo, 2016). Converging evidence further demonstrates that the hippocampus exhibits systematic visual responses, including sensitivity to the retinotopic position of visual stimuli (Knapen, 2021; Silson et al., 2021; Angeli et al., 2024) and egocentric viewing position (Gulli et al., 2020; Piza et al., 2024). Viewed through this lens, retinotopic sensitivity and egocentric view coding suggest that hippocampal representations are structured in formats well suited for predicting the sensory consequences of eye movements. Our findings extend this view by showing that hippocampal theta dynamics are temporally aligned with the moments at which such predictions would be most useful — immediately before and after visual sampling — thereby linking representational structure to the dynamics of active vision.

It is important to note several limitations of our work. First, comparisons between accurate recollection (true-positive trials) and the feeling of remembering (false-positive trials) were statistically underpowered because of high memory performance. However, preliminary analyses of such comparisons (TP and FP trials) revealed qualitatively similar results to the primary contrast (TP and TN trials) (Fig. S5), supporting our conclusion that memory-guided viewing displays unique power and coherence dynamics time-locked to fixation. Second, future work incorporating richer associative memory measures -- such as ratings of vividness or recollective confidence -- may help clarify how theta power and coherence scale with the quality of retrieved representations. A further limitation is that fixation-locked analyses were restricted to temporally isolated fixations, excluding fixations that occurred too closely together to separate their neural responses. This criterion improved the interpretability of the fixation-locked signal but reduced the number of analysed fixations and may prevent generalisation to denser periods of visual exploration. Future work should test whether similar effects are observed using approaches that model overlapping responses across the full fixation sequence.

A few open questions remain. Theta coherence is thought to facilitate hippocampal interactions with cortical and subcortical structures involved in memory retrieval and naturalistic perception (Solomon et al., 2017; Sonkusare et al., 2019; Zhang et al., 2021), yet our analyses focused on intra-hippocampal dynamics. A plausible next step will be to examine how theta synchrony across the hippocampus, visual cortex, and oculomotor regions supports memory-guided eye movements. Mechanistically, future studies are needed to establish, in a causal manner, the functional role of theta coherence during memory-guided saccades. Finally, the role of theta coherence during memory-guided behaviour in active visual contexts, where large-scale visual actions, such as head turns (Mynick et al., 2025) or body reorientations, complement eye movements, remains unexplored and may reveal deeper links between memory, attention, and active vision.

In conclusion, hippocampal theta dynamics are tightly coupled to visual sampling during naturalistic episodic retrieval. When memory guides viewing, broadband theta phase synchrony increases around fixation onset, whereas theta power is suppressed following fixation, relative to memory-independent viewing. This temporal dissociation suggests that theta supports memory-guided vision through complementary synchrony and power mechanisms, coordinating mnemonic signals with active visual exploration. Together, these findings position hippocampal oscillations as a central substrate through which prior knowledge shapes when, where, and how information is sampled in the natural visual environment.

## Acknowledgements

We are grateful to Kari L. Hoffman for insightful discussions.

